# Evaluation of Different PCR Assay Formats for Sensitive and Specific Detection of SARS-CoV-2 RNA

**DOI:** 10.1101/2020.06.24.168013

**Authors:** Jeremy Ratcliff, Dung Nguyen, Monique Andersson, Peter Simmonds

## Abstract

Accurate identification of individuals infected with SARS-CoV-2 is crucial for efforts to control the ongoing COVID-19 pandemic. Polymerase chain reaction (PCR)-based assays are the gold standard for detecting viral RNA in patient samples and are used extensively in clinical settings. Most currently used quantitative PCR (RT-qPCRs) rely upon real-time detection of PCR product using specialized laboratory equipment. To enable the application of PCR in resource-poor or non-specialist laboratories, we have developed and evaluated a nested PCR method for SARS-CoV-2 RNA using simple agarose gel electrophoresis for product detection. Using clinical samples tested by conventional qPCR methods and RNA transcripts of defined RNA copy number, the nested PCR based on the RdRP gene demonstrated high sensitivity and specificity for SARS-CoV-2 RNA detection in clinical samples, but showed variable and transcript length-dependent sensitivity for RNA transcripts. Samples and transcripts were further evaluated in an additional N protein real-time quantitative PCR assay. As determined by 50% endpoint detection, the sensitivities of three RT-qPCRs and nested PCR methods varied substantially depending on the transcript target with no method approaching single copy detection. Overall, these findings highlight the need for assay validation and optimization and demonstrate the inability to precisely compare viral quantification from different PCR methodologies without calibration.

## INTRODUCTION

SARS-Coronavirus-2 (SARS-CoV-2), a human-infective member of the *Betacoronavirus* genus (family *Coronaviridae*), was first identified in the Hubei Province of China in late 2019 as the causative agent behind an increased number of cases of respiratory illness occasionally leading to acute respiratory distress and death.^1-3^ The outbreak was declared a Public Health Emergency of International Concern by the World Health Organization on January 30^th^, 2020 and the associated disease was named COVID-19 on February 11^th^, 2020.^4,5^ The disease has since spread globally and by June 23^rd^ has infected nearly 9 million individuals in over 180 countries, causing at least 465,000 deaths.^6,7^

The ability to accurately identify and diagnose asymptomatically and symptomatically infected patients is crucial for efforts aimed at limiting person-to-person transmission and controlling the outbreak.^8-10^ The standard method for diagnosing viral infections is through the detection of viral nucleic acid in clinical samples. Reverse-transcriptase quantitative polymerase chain reaction (RT-qPCR) is the gold standard used in most diagnostic laboratories.^11^ Probe-based RT-qPCR relies on the binding and amplification of three oligonucleotides (two primers and one internal fluorescent probe) and the accumulation of fluorescence signal mediated by DNA polymerase activity. RT-qPCR is not widely accessible as the method relies upon the use of expensive real-time PCR platforms and the probe component of the assay is typically the most costly reagent. An alternative, more cost-effective diagnostic method for SARS-CoV-2 RNA is nested PCR. Nested PCR is based on the use of two sequential PCR amplifications wherein the secondary set of primers target sequences nested within the amplicon produced by the first round amplification. Compared to conventional PCR, which uses a single round of replication, nested PCR has increased sensitivity and decreased risk of amplification of non-specific products.^12^ Nested PCR methods were developed for SARS,^13^ but no nested PCR method for SARS-CoV-2 has yet been published.

In this study, a nested PCR assay for SARS-CoV-2 has been developed and its performance for an *in vitro* transcribed partial transcript of the RNA-Dependent RNA Polymerase (RdRP) gene and three transcripts containing the entire viral genome were compared against a corrected version of the RdRP-directed oligonucleotides from the Charité Institute of Virology (Charité)^14^ and the United States Centers for Disease Control and Prevention (CDC) N1 primer/probe set.^15^ Unexpectedly, the performances varied substantially depending on the detection method and target assayed, underpinning the need for in-house validation and optimization. The result also challenges the notion that Ct values presented without context could be an informative metric for the progression of disease and can be compared across different amplification techniques and laboratories.

## MATERIALS AND METHODS

### Primer design

1^st^ round nested PCR primers (nF1, nR1) were designed to anneal to conserved regions flanking a 1.1kb region of the RdRP gene of SARS-CoV-2. 2^nd^ round nested primers (nF2, nR2) were designed to anneal to conserved regions on the amplicon product of nF1 and nR1 amplification. Both sets of nested PCR primers were acquired from Integrated DNA Technologies. The Charité RT-PCR was based upon previously described primer/probes for the RdRP gene^14^ but with modifications to the antisense primer to ensure complete sequence complementarity with SARS-CoV-2 sequences (Table 1; differences underlined). All primers and probes for the Charité and CDC N1 PCRs were obtained from ATDBio. All primer sequences and working concentration are available in Table 1.

**Table 1:**
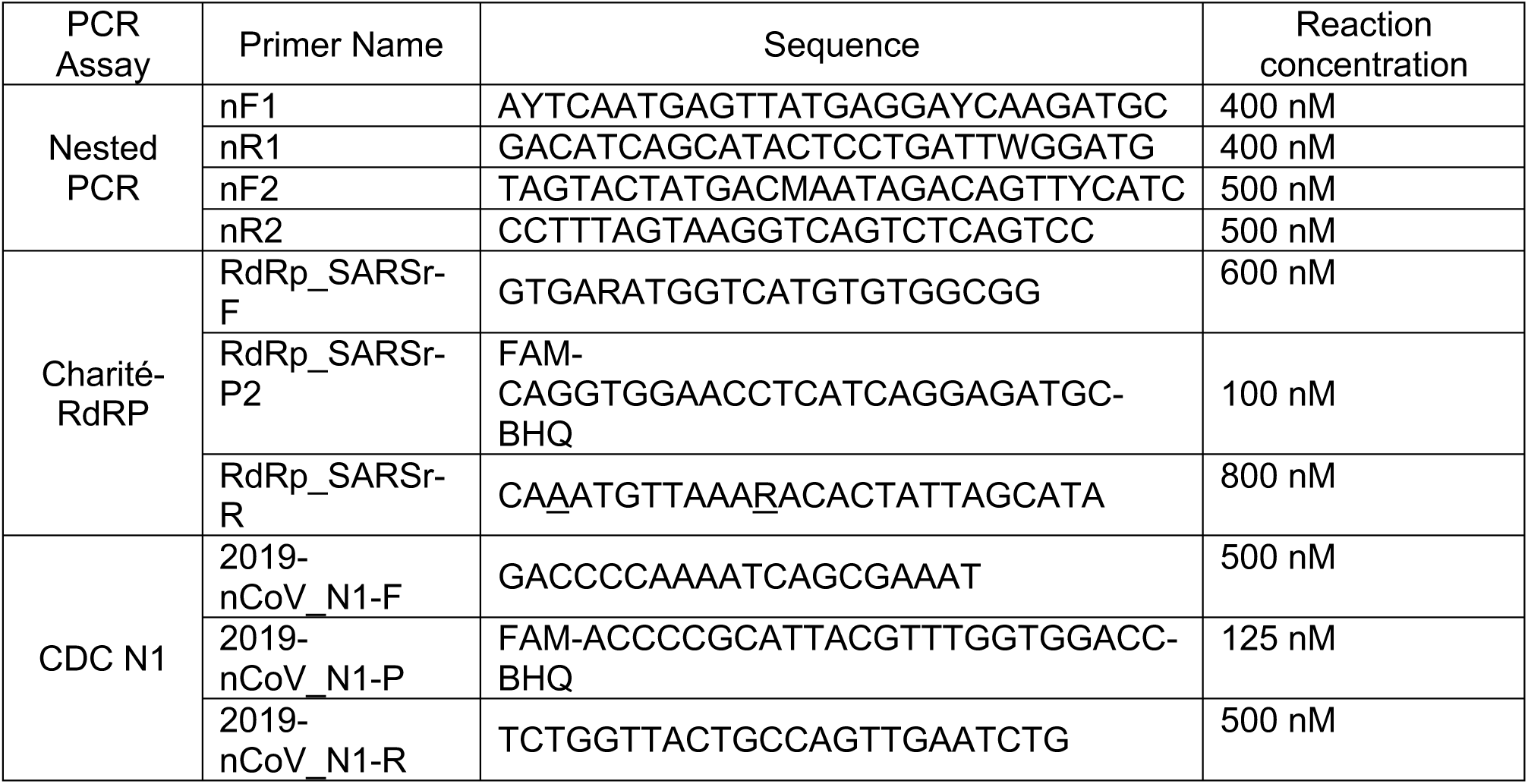
Primer and Probe Sequences for Nested PCR and RT-qPCR.

**Table 2:**
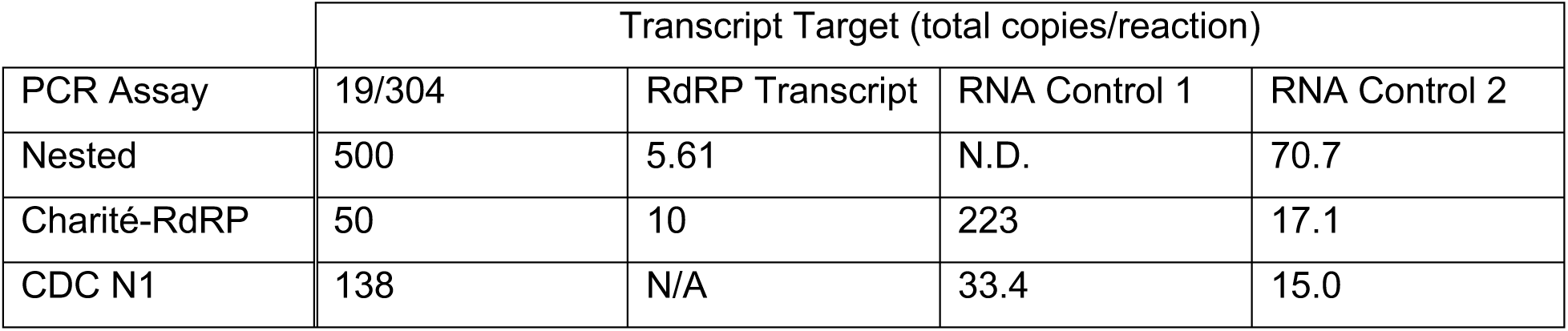
50% endpoints for PCR/target pairs. 50% endpoints of detection for serial dilutions of four SARS-CoV-2 transcripts by three PCR assays. Nested PCR is detection rate over five replicates; Charité-RdRP and CDC N1 are detection rate over eight replicates. 50% endpoints were calculated by the Reed-Muench method.

### Clinical Samples

Residual RNA from clinical samples was eluted in buffer AVE (Qiagen) following extraction using a Qiasymphony DSP virus/pathogen minikit. SARS-CoV-2 infection status was determined by RT-qPCR operated on the Rotor-Gene Q using an RdRP gene target validated by Public Health England (PHE)^14^ or the Altona RealStar SARS-CoV-2 RT-PCR Kit targeting the E and S genes.

### Nested PCR

The nested PCR uses two sequential PCR amplifications for highly specific and sensitive amplification of target sequences (primer sequences and concentrations listed in Table 1). For the first round of amplification, a 25 µL reaction mix containing 5 µL of RNA extract, 12.5 µL of 2X Quantitect Probe RT-PCR Master Mix (Qiagen), 0.5 µL of Superscript III Reverse Transcriptase (ThermoFisher Scientific), 5 µL of 5X 1^st^-round primer mix, and 2 µL of PCR-grade water was amplified on a thermal cycler using the following settings: reverse transcription at 50°C for 30 m, activation at 95°C for 15 m, 40 cycles of 95°C for 15 s, 55°C for 30 s, and 68°C for 1 m, and a final extension of 68°C for 5 m. For the second round amplification, a 25 µL reaction mix containing 1 µL of the first round product, 5 µL of 5X GoTaq Green Master Mix (Promega), 0.125 µL of 5u/µL GoTaq G2 polymerase (Promega), 2 µL of 2.5 mM dNTP mix (Stratech Scientific), 5 µL of 2^nd^-round 5X primer mix, and 11.875 µL of PCR-grade water was amplified on a thermal cycler using the following settings: 95°C for 5 m, 40 cycles of 95°C for 30 s, 55°C for 30 s, and 72°C for 1 m, and a final extension of 72°C for 5 m. The presence of SARS-CoV-2 RNA was confirmed through UV visualization of a PCR product of the expected length on a 2% agarose gel stained with Sybr-Safe DNA Gel Stain (ThermoFisher Scientific).

### Real-time Reverse Transcription PCR

Master Mix contents and PCR system settings were standardized for both primer/probe concentrations. A 20 µL reaction mix containing 12.5 µL of 2X Quantitect Probe RT-PCR Master Mix (Qiagen), 0.5 µL of RT mix from the kit, 5 µL of 5X oligo mix, and 2 µL of PCR-grade water was mixed with 5 µL of RNA extract in a MicroAmp Fast Optical 96-well reaction plate (ThermoFisher Scientific). Amplification was performed on an Applied Biosystems StepOnePlus Real-Time PCR System (ThermoFisher Scientific) with the following settings: 50°C for 30 m, 95°C for 15 m, and 40 cycles of 94°C for 15 s and 60°C for 1 m.

### Transcripts

A 1.1kb transcript of the RdRP was synthesized from the first round nested PCR product of the COV02 clinical sample. The fragment was ligated with the pJet1.2 plasmid using the CloneJET PCR Cloning Kit (ThermoFisher Scientific). The recombinant plasmid was linearized using the Ncol restriction enzyme (ThermoFisher Scientific) and the MEGAscript® T7 Kit (ThermoFisher Scientific) was used for *in vitro* transcription of the fragment. NIBSC research reagent 19/304 containing packaged SARS-CoV-2 RNA was the kind gift of Giada Mattiuzzo. 19/304 was extracted using the Viral RNA Mini Kit (Qiagen) and eluted into 50 µL of buffer AVE. Synthetic SARS-CoV-2 RNA Control 1 - MT007544.1 and Control 2 - MN908947.3 were synthesized by Twist Bioscience. The accession numbers for RNA control 1 and RNA control 2 have no nucleotide differences in the binding locations of oligonucleotides used in this study.

### Sensitivity of PCR-based assays

Experiments to determine the sensitivity of the two RT-qPCR methods and nested PCR were completed using serial dilutions of each transcript (5*10^3^ to 10^−1^ copies/5 µL) in a previously described RNA storage buffer containing RNA storage solution (Thermo Fisher Scientific; 1 mM sodium citrate, pH 6.4), herring sperm carrier RNA (50 µg/mL), and RNasin (New England BioLabs UK, 100 U/mL).^16^ For nested PCR experiments, detection rate was assessed over five replicates using the methods described above and a positive result was the presence of a PCR product of the expected length. For RT-qPCR experiments, detection rate was assessed over eight replicate experiments using the methods described above and a positive result was an increase in signal that crossed the threshold value calculated by the machine for each experiment. The CDC N1 method had a cutoff value of Ct 35.

### Statistical Analysis

50% endpoints were estimated using the Reed-Muench method.^17^ Probit models were constructed using function glm in R version 3.6.2.^18^

## RESULTS

### Detection of Clinical Samples via Nested PCR

To determine the ability of the nested PCR to detect and differentiate SARS-CoV-2 RNA in extracts from clinical samples, 43 samples of known SARS-CoV-2 status were assessed. The nested PCR successfully produced a PCR product of the expected size in 33/35 positive samples (sensitivity of 94.2%) and a negative signal in 8/8 negative samples (specificity of 100%) (Figure 1, Figure S1). The nested PCR detected SARS-CoV-2 RNA from samples with a range of Ct values – from a minimum of 17.11 to a maximum of 36.57. The two false negative samples had relatively high Ct values of 29.6 and 34.66 but were not the most extreme targeted as demonstrated in Figure 1.

**Figure 1:**
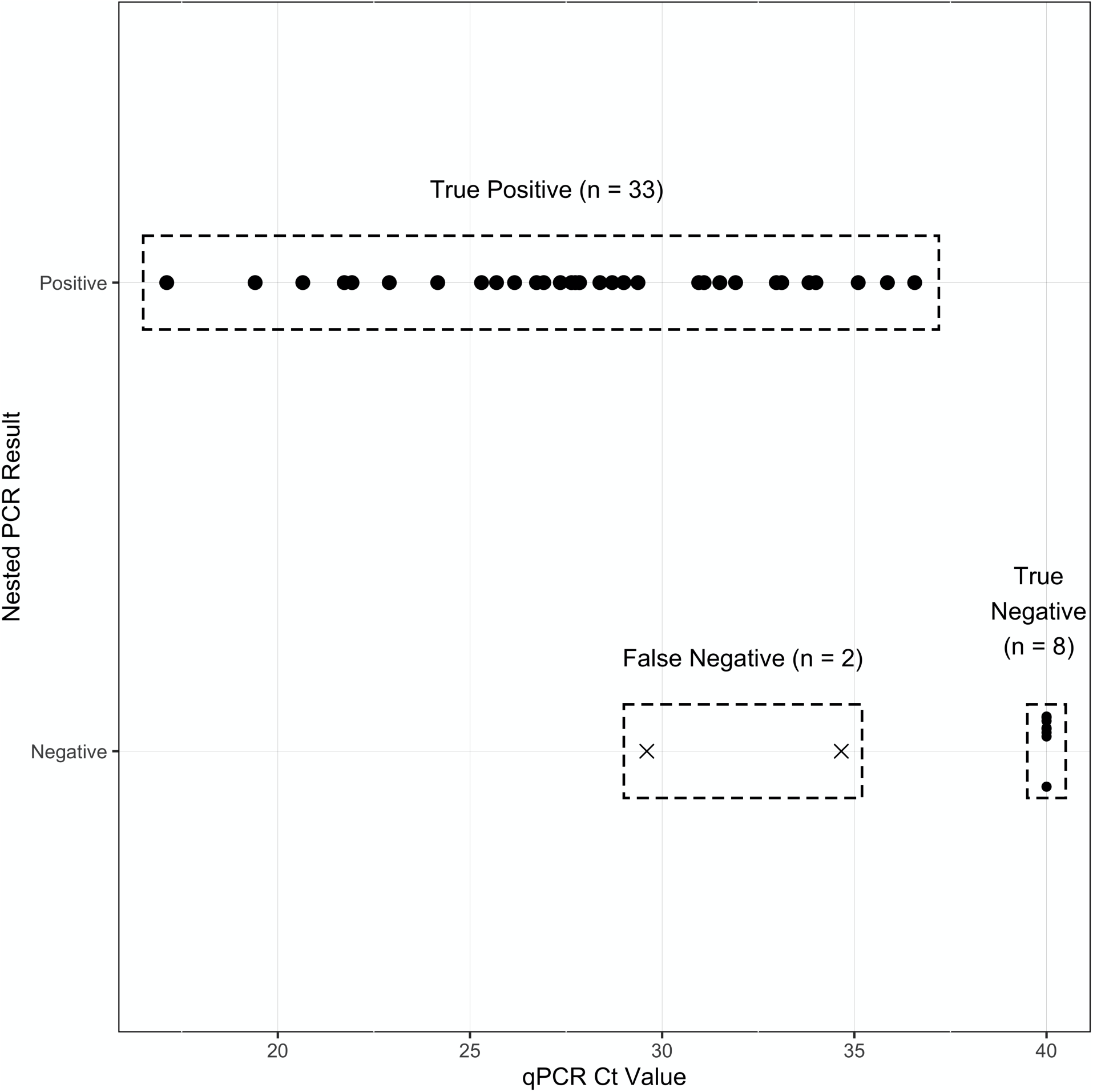
Performance of Nested PCR for clinical samples. Comparison of detection of SARS-CoV-2 RNA in clinical samples by nested PCR versus qPCR readout from the microbiology unit of the John Radcliffe Hospital. Samples negative via qPCR were given a representative Ct value of 40.

### Comparative Sensitivity

The sensitivity of the nested PCR and two RT-qPCR methods was compared by measuring the 50% endpoints (50EP) of detection for serial dilutions of the four transcripts described above (Table 1, Figure 2). The RdRP transcript does not contain the target sequences of the CDC N1 method and thus was not assessed using this method. No method proved to consistently be the most sensitive for all targets, and each method was the most sensitive for at least one target. The RdRP transcript was detected consistently by the nested PCR with a 50EP of 5.61 copies, making this the most sensitive for detecting the RdRp transcript. The Charité-RdRP RT-qPCR method was the most sensitive for the 19/304 transcript with a 50EP of 50 copies. The CDC N1 method was the most sensitive for both RNA Control 1 and RNA Control 2 with 50EPs of 33.4 and 15.0 copies, respectively. The differences between methods and a single target did not follow a clear trend. Between the CDC N1 and Charité-RdRP RT-qPCR methods, the CDC N1 method showed clear superiority for the RNA control 1 target (33.4 copies versus 223 copies) but only marginal superiority for the RNA control 2 target (15.0 copies versus 17.1 copies). Strikingly, the nested PCR was not able to detect RNA control 1 with any appreciable regularity even at the highest copy number tested despite robust detection of RNA control 2.

**Figure 2:**
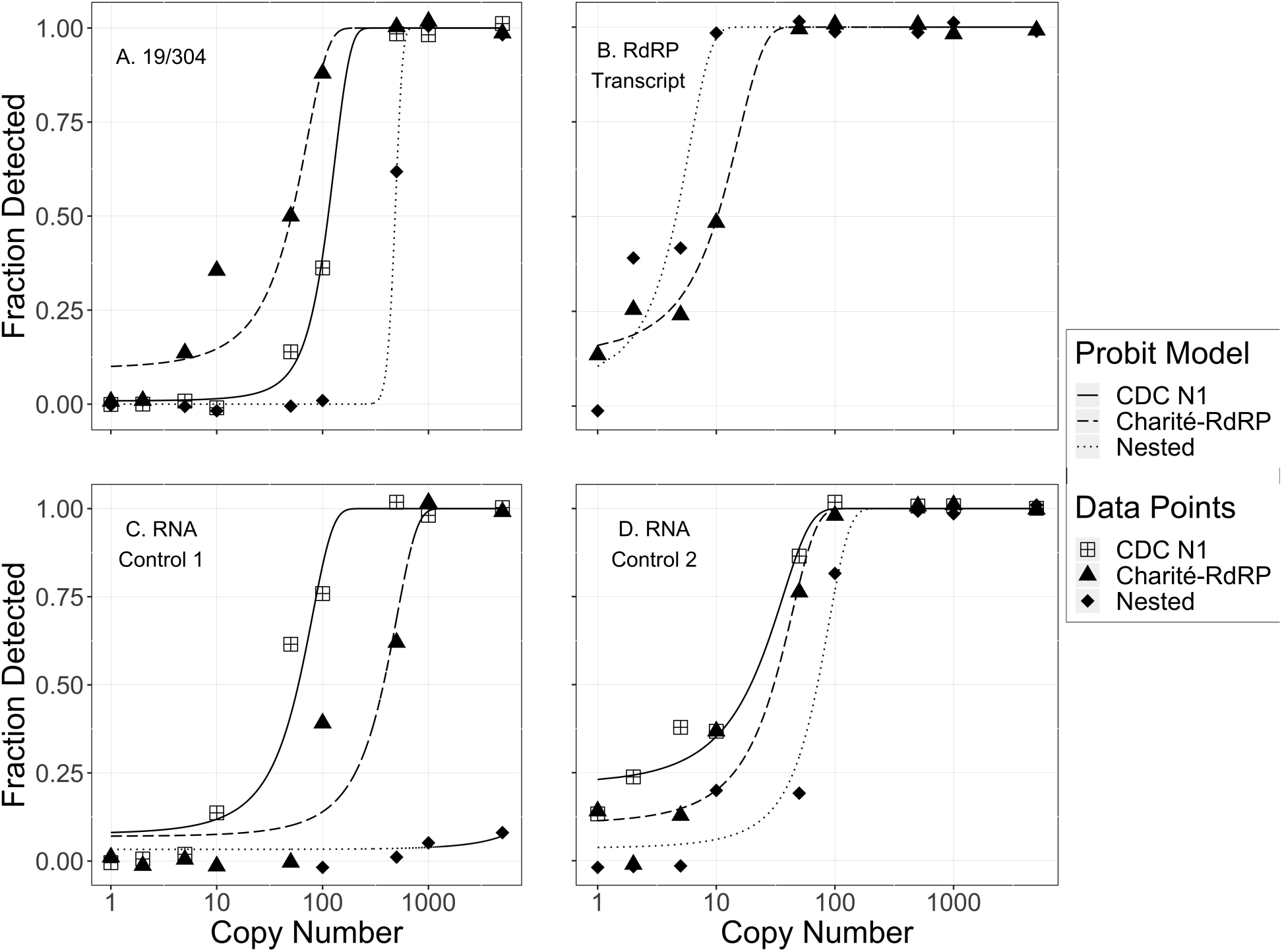
Comparative detection performance for nested PCR and Charité-RdRP and CDC N1 qPCR. Comparative performance of detection methods for four targets. Dots for qPCR experiments represent fraction detected of eight replicate reactions. Dots for nested PCR experiments represent fraction detected of five replicate reactions (15 replicates for RNA control 1). Lines represent probit regression models.

## DISCUSSION

This study establishes the first published nested PCR method for the sensitive detection of SARS-CoV-2 in RNA extracted from nasopharyngeal swab samples. The nested PCR showed highly concordant results with RT-qPCR performed at a hospital diagnostic laboratory, with a sensitivity of 94.2%, specificity of 100%, and overall agreement of 95.3%. The use of nested PCR is a practical alternative for laboratories without real-time PCR systems, but does require an RNA extraction step. Whilst there has been a global shortage of necessary reagents,^19^ an increasing number of novel extractions methods have been described in the literature.^20-23^ Furthermore, nested PCR is also less amenable to automation and requires equipment for casting agarose gels and amplicon visualization using UV light. The poor performance of the nested PCR for detecting RNA control 1 is hypothesized to be due to the design of the transcript standard. Twist Bioscience synthetic SARS-CoV-2 transcripts are six, non-overlapping 5 kb fragments. The drastically low sensitivity but presence of PCR products of the correct length strongly suggests that the fragmentation process separates the nF1 and nF2, nR1, and nR2 binding locations onto two different fragments. This result underscores the notion that not all transcript standards are equal for different PCR-based detection modalities.

Secondarily, this study demonstrates inconsistent sensitivity for SARS-CoV-2 targets for different PCR-based detection modalities. The results of this study point to remarkably different performances between the three PCR methods analyzed for different SARS-CoV-2 RNA transcripts. This inconsistently between different RT-qPCR methods has been noted by other authors. In presenting the Charité-RdRP primer/probe set, Corman et al. found slight differences between their E, N, and RdRP primer/probe sets, with the RdRP set performing the best with a 95% detection probability of 3.8 copies/reaction for SARS-CoV-2 genomic RNA.^14^ Corman et al. also reported a 95% detection probability of 5.2 copies/reaction for the E gene assay – this is in conflict with a 95% detection probability of 100 copies/reaction reported by Institut Pasteur using the same primer/probe set.^24^ Igloi et al. investigated the performance of 13 commercial RT-qPCR kits and found variable performance between the RT-qPCR kits, with several kits having 10 fold differences in sensitivity for different gene targets and one kit having a 2 log difference in sensitivity between their E and RdRP/N preparations.^25^ Using *in vitro* transcribed small transcripts of five SARS-CoV-2 genes, Vogels et al. evaluated nine primer/probe sets, including the Charité-RdRP and CDC N1 sets.^26^ Vogels et al. found that eight of the nine primer/probe sets had lower limits of detection of 500 copies/reaction, and that the Charité RdRP set was unable to detect any replicates with 500 copies/reaction, although they did alter the primer and probe concentration from those suggested by Corman et al. Lastly, Kudo et al. designed a multiplex RT-qPCR using the CDC N1, N2, and RNAse P primer and probe sets.^27^ Using a full-length control and reanalyzing the raw data using the Reed-Muench method, the multiple RT-qPCR demonstrated a 50EP of 30.0 copies/reaction for the CDC N1 primers and 107.7 copies/reaction for the CDC N2 primers.

These results are compatible with anecdotal evidence from several labs in the United Kingdom and elsewhere that consistent low level detection is unreliable and aberrant results are common. This broader experience suggests that the issue may lie within the generalized approach of detecting SARS-CoV-2 RNA by PCR rather than being specific to individual assays. One possible factor limiting the sensitivity of PCR assays for SARS-CoV-2 may originate from the highly structured nature of the genome as measured by standard thermodynamic RNA structure prediction methods.^28^ SARS-CoV-2, as well as other coronaviruses, has extensive RNA secondary structure elements peppered throughout its genome – approximately 62% - 67% of bases may be involved in base pairing. It is conceivable that, if a section of the target sequence is embedded within highly energetically favoured RNA secondary structure, binding of the PCR oligonucleotides could be competitively inhibited, delaying the initiation of the PCR reaction. This effect could explain the unusual findings that no PCR based assay for SARS-CoV-2, whether in this study or others, have been able to achieve single copy detection sensitivities.

Understanding the association between viral load and disease progression is important for the selection of therapies. Tom et al.^29^ recently proposed reporting RT-qPCR Ct values rather than binary PCR test results within clinical reports as a means of estimating the viral load of patients due to the inverse correlation of Ct value with copy number. While informative when tracking the progression of Ct values for a single patient over time or comparing to other patients within the same diagnostic pipeline, the results of this study do not support making direct comparisons between data gained from different laboratories as Ct values are not readily comparable. For these values to be informative, Ct values published in the literature should contain the methods used as well as estimates of in-house correlation with copy. Alternatively, as also recommend by Tom et al., reports could provide the Ct value as binned within ranges also seen in the laboratory (i.e. in the top quartile of Ct values seen in this laboratory).

This current study is limited in that the standardized thermal cycler temperatures and concentrations of Mg^2+^ recommended by the Quantitech RT-PCR kit were used for both the Charité-RdRP and CDC N1 primers. It is feasible that slight alterations to the annealing temperature and salt concentration could result in marginal increases in sensitivity, but these changes would have to be assessed and calibrated on a lab by lab basis and may have different values depending on the RT-qPCR master mix and PCR system used.

## ACKNOWLEDGEMENTS

The authors acknowledge the staff of the Microbiology Department of the John Radcliffe Hospital for their work in processing and testing clinical samples. The authors acknowledge Dr. Alexander Mentzer for his assistance in procuring reagents for RT-qPCR and Dr. John Taylor for his gift of RNA control 1 and RNA control 2. The authors want to especially thank Professor Tom Brown and the team at ATDBio for rapid synthesis and delivery of oligonucleotides.

## CONFLICTS OF INTEREST

The authors report no conflicts of interest.

## Supplementary Material

**Figure S1:**
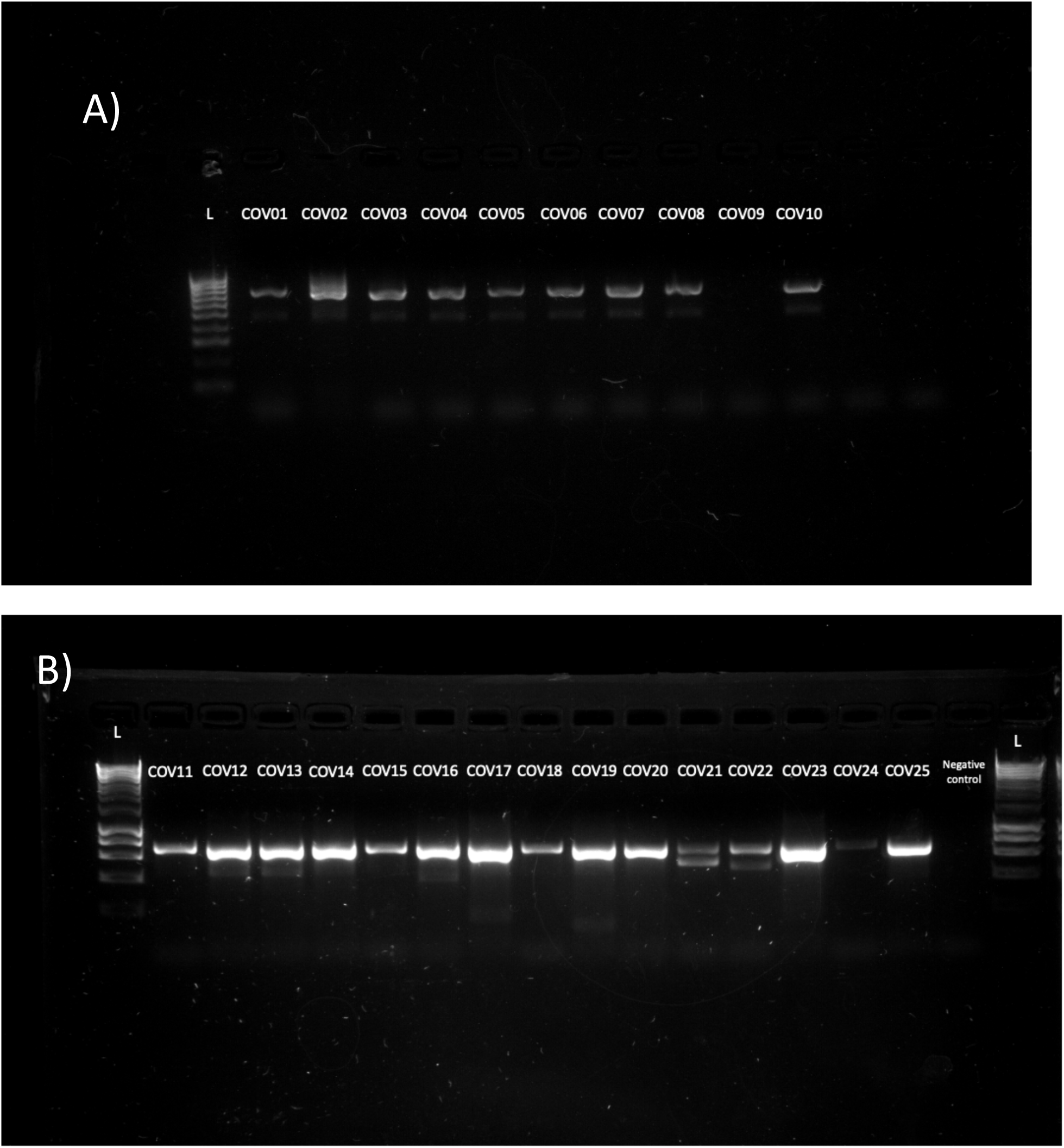
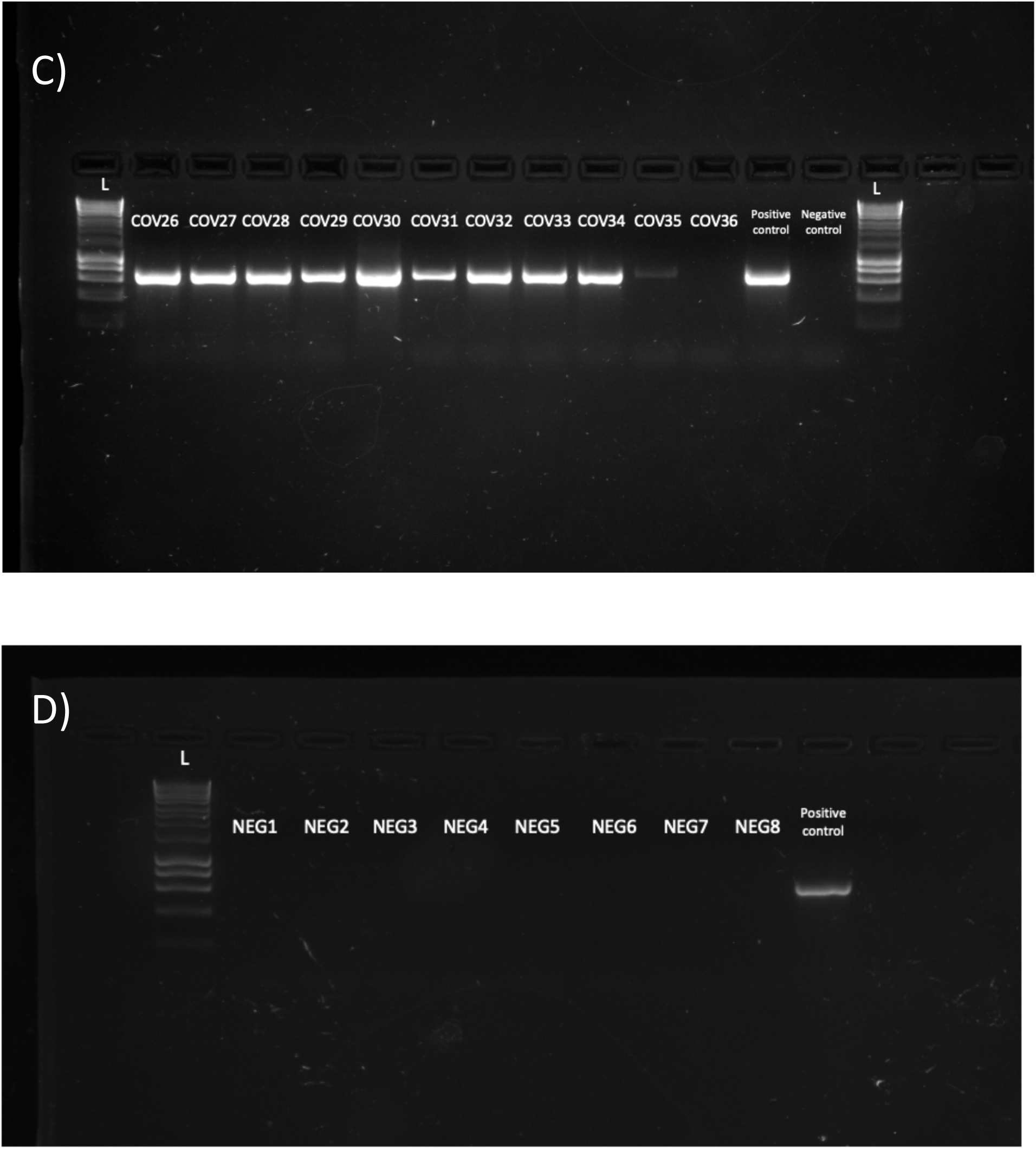
Nested PCR amplification of clinical samples. Nested PCR detection of clinical samples assessed at the John Radcliffe Hospital via RT-qPCR. COV1-COV36 (panels A-C) were positive samples, NEG1-NEG8 (Panel D) were negative samples. Panel A has a 100 bp molecular ladder (New England Biolabs); Panels B-D have 1kb molecular ladder (New England Biolabs).

## Notes

### Competing Interest Statement

The authors have declared no competing interest.

### Summary of Updates

- Update spelling mistake in author list - Update clarification in methods

